# DiffVar: A new method for detecting differential variability with application to methylation in cancer and aging

**DOI:** 10.1101/008847

**Authors:** Belinda Phipson, Alicia Oshlack

## Abstract

Methylation of DNA is known to be essential to development and dramatically altered in cancers. The Illumina HumanMethylation450 BeadChip has been used extensively as a cost-effective way to profile nearly half a million CpG sites across the human genome. Here we present DiffVar, a novel method to test for differential variability between sample groups. DiffVar employs an empirical Bayes model framework that can take into account any experimental design and is robust to outliers. We applied DiffVar to several datasets from The Cancer Genome Atlas, as well as an aging dataset. DiffVar is available in the *missMethyl* Bioconductor R package.

## BACKGROUND

DNA methylation is crucial for normal embryonic development with roughly 3-6% of all cytosines methylated in normal human DNA [1]. However, methylation changes are known to accumulate with age [2], with up to 30% of CpG sites changing methylation status within the first 1.5 years of life [3]. In addition, aberrant methylation patterning is associated with many diseases. In particular, in cancer cells, disruption of normal methylation events are very common with the number of genes undergoing CpG island promoter hypermethylation increasing during tumour development, combined with an extensive loss of DNA methylation in other genomic regions [1, 4]. This phenomenon is not consistent across all cancers however; distinct DNA methylation patterns have been observed between sub-types of the same cancer [5–7]. Akalin et al. [5] show that one sub-type of Acute Myeloid Leukemia shows widespread hypermethylation in promoter regions and CpG islands neighbouring the transcription start site of genes, while a second sub-type displayed extensive loss of methylation at an almost mutually exclusive set of CpGs in introns and intergenic CpG islands and shores. Abnormal methylation events can potentially silence tumour suppressor or growth regulatory genes, activating novel pathways that contribute to tumour progression [8]. However, epigenetic changes are potentially reversible, as it is possible to reactivate genes that have been silenced by methylation [1], making them an attractive therapeutic target. Hence the study of DNA methylation in cancer remains an important topic of interest with much still to be discovered.

Cancer is a heterogeneous disease. Within each type of cancer there is the potential for tumour growth to be driven by perturbations in many different molecular pathways, and these perturbations will vary from individual to individual. Epigenetic instability or the loss of epigenetic control of important genomic domains can lead to increased methylation variability in cancer, which may contribute to tumour heterogeneity [9]. In a large study profiling 1505 CpG sites of 1628 human samples, of which 1054 were tumours, Fernandez et al. [10] observed little variation in the DNA methylation patterns of normal tissue, while the established tumours showed greater CpG methylation heterogeneity. Hansen et al. [9] propose that increased epigenetic heterogeneity in cancer could underlie the ability of cancer cells to adapt rapidly to changing environments. Hence studying the heterogeneity of cancers could lead to better understanding of tumourigenesis.

As mentioned, it has been shown that DNA methylation patterns are associated with an individuals’ age [2]. Interestingly, both tumour development and aging are processes which result in the global loss of genome wide DNA methylation combined with gains in CpG island promoter methylation [11, 12]. Hence it has been speculated that the accumulation of epigenetic alterations during aging might contribute to tumourigenesis [12]. Several technologies are available for profiling DNA methylation, both array and sequencing based. While the cost of next generation sequencing has dramatically decreased, it is still too expensive for many large studies to profile widespread methylation. The introduction of the Illumina 450K human methylation array is a more affordable alternative for measuring genome-wide DNA methylation at 482,421 CpG sites. Consequently, thousands of tumour and normal tissues have been profiled using these arrays, with numerous cancer datasets now publicly available through The Cancer Genome Atlas (TCGA) [13, 14]. Similarly, these arrays are being used to profile methylation differences between disease cases and normal controls in so called eigenome-wide association studies (EWAS) [15].

To date, the main focus when analysing DNA methylation data has been detecting CpG sites that are differentially methylated between groups. Methods to detect differences in means for high-dimensional biological data are well-established and include approaches such as those taken in the Bioconductor [16] software packages *limma* [17]*, minfi* [18]*, edgeR* [19] and *DESeq* [20]. A CpG site that is statistically significantly differentially methylated between groups (for example, cancer versus normal) will have different group means; however, the measurements within each group will tend to be quite consistent (Figure 1A). Recently, several papers have observed consistent methylation between normal samples and highly variable methylation between cancer samples, arguing that identifying features that differ in terms of variability may be just as relevant or important as differential methylation for understanding disease phenotypes [21–25]. In other words, there is interest in identifying differentially variable CpG sites, where the samples in one group have consistent methylation values and the samples in the other group have highly variable methylation values (Figure 1B).

**Figure 1.**
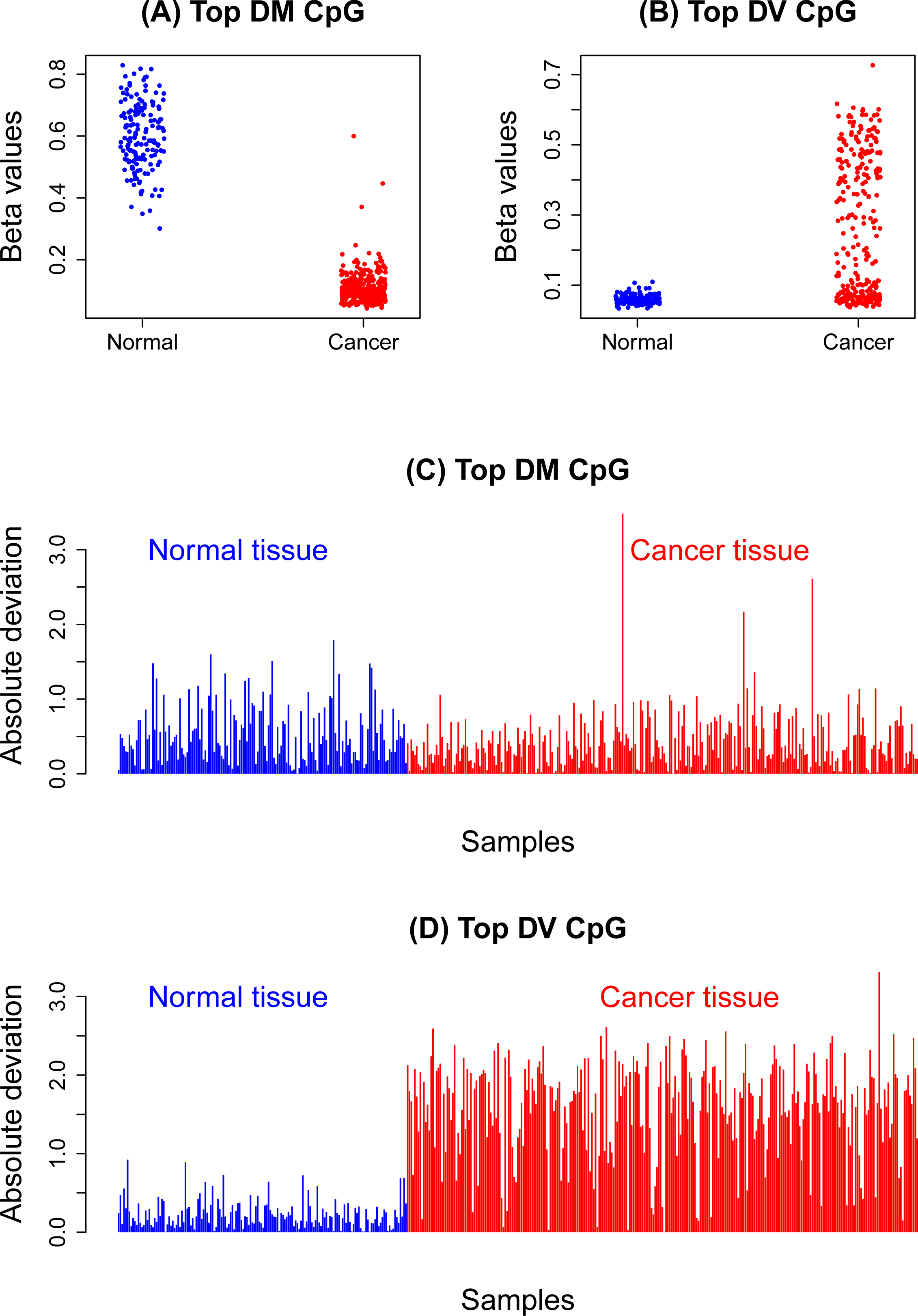
Differential methylation (DM) and differential variability (DV) in the kidney cancer methylation dataset. The plots in (A) and (C) show the M values and absolute deviations of the most significant differentially methylated CpG site between the normal and cancer samples. There is an obvious shift in mean between cancer and normal (A), but from (C) the variability in both groups looks very similar. The plots in (B) and (D) show the M values and absolute deviations of the most significant differentially variable CpG site between the normal and cancer samples. It is clear that the variability of the cancer and normal groups are very different, with very large deviations in the cancer group and consistently small deviations in the normal group apparent in (D).

Methods for detecting differential variability in high dimensional ‘omics data have not been well addressed in the literature to date. Jaffe et al. [24] have a sophisticated method to detect differential variability over regions, specifically developed for the CHARM array [26]. Bar et al. [27, 28] propose a three component mixture model on the ratio of the sample variances from treatment and control groups. The method aims to separate features that are not differentially variable, have inflated variance in the treatment group, or inflated variance in the control group. In addition, they perform empirical Bayes shrinkage on the inflation factors to stabilise the estimates. Other attempts for determining CpG site-wise differential variability include the F-test [9, 29] and the Bartlett test [25]. Unfortunately, the F-test and Bartlett test are known to be highly sensitive to outliers [30]. While Hansen et al. [9] do not address the issue of outliers in the data when using an F-test for equality of variances; Ho et al [29] perform an outlier removal step prior to testing for differential variability. By contrast, Teschendorff and Widschwendter [25] implement the Bartlett test and claim that features that are differentially variable due to outliers are of interest. Their application is specific to the early stages of carcinogenesis, where they hypothesised that outliers may play an important role. It has been observed, however, that outliers are often a result of technical and biological artefacts rather than being biologically relevant to disease. For example, a technical artefact could arise due to processes surrounding the technology and biological artefacts could include stromal contamination of a tumour sample. Microarrays can suffer from spatial artefacts [31] and outliers arising from sample specific GC content biases have been reported in RNA sequencing data [32]. Mislabelled samples can also lead to outlying observations [33]. Hence, a method that successfully identifies differentially variable sites with a broader distribution of methylation values, such as that in Figure 1B, is desirable. Furthermore, the F-test and Bartlett test assume that the data is normally distributed, which is not the case for methylation data [34].

Here we present a new method for detecting differential variability for individual CpG sites in methylation data. Our approach is inspired by Levene’s z-test [35]. It is a simple and computationally efficient test that is robust against non-normality and outliers. A major advantage of our method is that it is suitable for any experimental design; it is not limited to a two-group scenario. The method, called *DiffVar*, is available as a function in the *missMethyl* R package available from Bioconductor [36], and depends on the *limma* framework. We applied DiffVar to several publicly available cancer datasets from TCGA, as well as a publicly available aging dataset [2]. When we applied DiffVar to the cancer data sets from TCGA we observed that a large proportion of the top differentially variable CpG sites are found in CpG islands. Interestingly, the top differentially variable CpG islands tend to differ from cancer to cancer. We further found that the 10,000 top ranked differentially variable CpG sites have very little overlap with the 10,000 top ranked differentially methylated CpG sites, consistent with the findings in Teschendorff and Widschwendter [25]. Applying DiffVar to an aging dataset revealed that the centenarians have highly variable methylation compared to newborns and approximately 17% of the differentially variable CpGs were also differentially methylated.

## RESULTS

### DiffVar: A new method to identify differentially variable features

The focus of this paper is on methylation data from Illumina’s Infinium HumanMethylation450 BeadChip, although our method can be applied to any high dimensional data such as gene expression data. The output from the array consists of two measurements for each CpG site, the methylated and unmethylated intensity signal. These two measurements can be used to calculate either *β* values, which capture the proportion of methylation at each CpG site, or M values, defined as the log_2_-ratio of the methylated to unmethylated intensity. Since M values do not display the severe heteroscedasticity that occur with *β* values [34], DiffVar is performed on M values. If only *β* values are available they can be transformed to M values using a logit transformation, M = logit(*β*). Similarly M values can be converted back to *β* values for interpretation and visualisation. For more details on the array and raw data see ‘Materials and Methods’.

As mentioned, our method is inspired by Levene’s z-test [35]. Intuitively, a measure of variability can be thought of as the distance of each point within a group from the group mean. Figures 1C and 1D show examples of the CpG sites from Figure 1A and 1B across a large kidney cancer and normal dataset from TCGA (see ‘Materials and Methods’). Each sample is plotted as a bar with the height of the bar equal to the absolute deviation from the group mean. Highly variable groups will be characterised by consistently large deviations (Figure 1D, red bars representing the cancer group) and low variability groups will have consistently small deviations (closer to zero) about the group mean (Figure 1D, blue bars representing the normal group). In order to determine if one group is more variable than another, we can simply perform a t-test on the absolute or squared deviations of the M values from the group mean. This tests the null hypothesis that the group variances are equal. CpG sites that have deviations consistently larger in one group compared to another group will be identified as significantly differentially variable. Figure 1C shows the absolute deviations from the group means for the top differentially methylated CpG site. Although there is a shift in mean M values between the two groups, the variations about the mean are similar in both groups; hence this CpG site will not be significantly differentially variable.

In high dimensional data, it is well-known that a simple t-test can result in many false discoveries, hence we employ an empirical Bayes modelling framework to stabilise the t-statistics [37]. Additional variables, which may influence the variability within groups, can be taken into account within the linear modelling framework. This ensures that any experiment that can be summarised by a design matrix can be accommodated. Unequal sample sizes are taken into account by multiplying the absolute or squared deviations from the group mean by a leverage factor, *n_k_ /(n_k_ +* 1), where *n_k_* is the sample size for group *k*. Moderated t statistics are computed and Benjamini and Hochberg false discovery rates (FDR) [38] are reported. In this manner, a list of differentially variable CpG sites is obtained. For a more formal definition of the statistical model see the Supplementary text.

### Simulation study

#### DiffVar correctly controls Type I error rate

Two strategies were used to determine type I error rate control. The first strategy generated M values under a hierarchical model while the second strategy involved resampling data from the kidney cancer data set, using only the non-diseased samples. Details of the simulations are provided in ‘Materials and Methods’. Briefly, a two group scenario with a sample size of 50 in each group was simulated. M values were generated for ten thousand CpG sites and with the same variance for each group. CpG sites were tested for differential variability using the F-test, Bartlett’s test, DiffVar with absolute deviations, and DiffVar with squared deviations. The numbers of significant raw p-values at cut-offs of 0.001, 0.01, 0.05 and 0.1 were counted for each of 1000 simulations. Table 1 summarises the results. Under the hierarchical model, the type I error rates are very close to the nominal p-value for all tests, showing that all tests have good type I error rate control in this scenario, although DiffVar with squared deviations appears conservative at 0.001. For the resampled data, the Bartlett test and the F test lose type I error rate control for all nominal p-values (Table 1). This is likely due to real data containing technical and biological artefacts that produce outliers. By contrast, DiffVar with absolute and squared deviations maintains type I error rate control over all nominal p-values. DiffVar with squared deviations is conservative at nominal p-values of 0.001 and 0.01.

**Table 1:** **Comparing the type I error rate for the four different methods**. Data was generated in two ways: under a hierarchical model, and by randomly selecting the non-diseased kidney samples. Median type I error rates are reported for 1000 simulations with no differentially variable or differentially methylated features. For simulations generated under the hierarchical model, the standard deviation with which the error rate is estimated ranges from approximately 0.00024 for rates near 0.001 to 0.0029 for rates near 0.1 with no notable difference between the methods. For the resampled data, the standard deviation ranges from approximately 0.0026 for rates near 0.001 to 0.026 for rates near 0.1 with DiffVar (sq) being noticeably less variable than the other methods.

| **Data generated under hierarchical model** |  |  |  |  |
| --- | --- | --- | --- | --- |
|  | **Nominal P-value** |  |  |  |
| **Test method** | **0.001** | **0.01** | **0.05** | **0.1** |
| Bartlett | 0.001032 | 0.01014 | 0.0504 | 0.1006 |
| F test | 0.001011 | 0.01003 | 0.05001 | 0.1001 |
| DiffVar (abs) | 0.0010029 | 0.01023 | 0.05094 | 0.1036 |
| DiffVar (sq) | 0.0007118 | 0.009800 | 0.0499 | 0.1018 |
| **Resampled data from kidney non-diseased samples** |  |  |  |  |
|  | **Nominal P-value** |  |  |  |
| **Test method** | **0.001** | **0.01** | **0.05** | **0.1** |
| Bartlett | 0.009 | 0.0257 | 0.07715 | 0.1331 |
| F test | 0.0089 | 0.0253 | 0.0763 | 0.1319 |
| DiffVar (abs) | 0.001 | 0.0104 | 0.0523 | 0.105 |
| DiffVar (sq) | 0.0004 | 0.00665 | 0.0448 | 0.0981 |

#### DiffVar is robust to outliers

To further explore the effect of outliers, the simulations under the hierarchical model were modified to incorporate 200 CpGs with a single outlier by randomly selecting one sample and substituting the maximum M value over all ten thousand simulated M values. From Table 2 it is again apparent that the Bartlett and F-test are extremely sensitive to outliers and have lost all type I error rate control, particularly for smaller nominal p-values. DiffVar with absolute or squared deviations is robust to outliers and maintains type I error control, although DiffVar with squared deviations is once again conservative at nominal rates of 0.001 and 0.01. Figure 2A shows the number of CpGs with outliers in the top 500 features ranked by each method. Although the F and Bartlett tests produce different p-values, they have similar ranking by p-value, resulting in overlapping curves for Figure 2A and 2B. It is immediately apparent that the top ranked features by Bartlett and F p-values are produced by probes with outliers. In contrast, DiffVar using either squared or absolute deviations does not preferentially rank the CpG sites containing outliers near the top of the list (Figure 2A).

**Figure 2.**
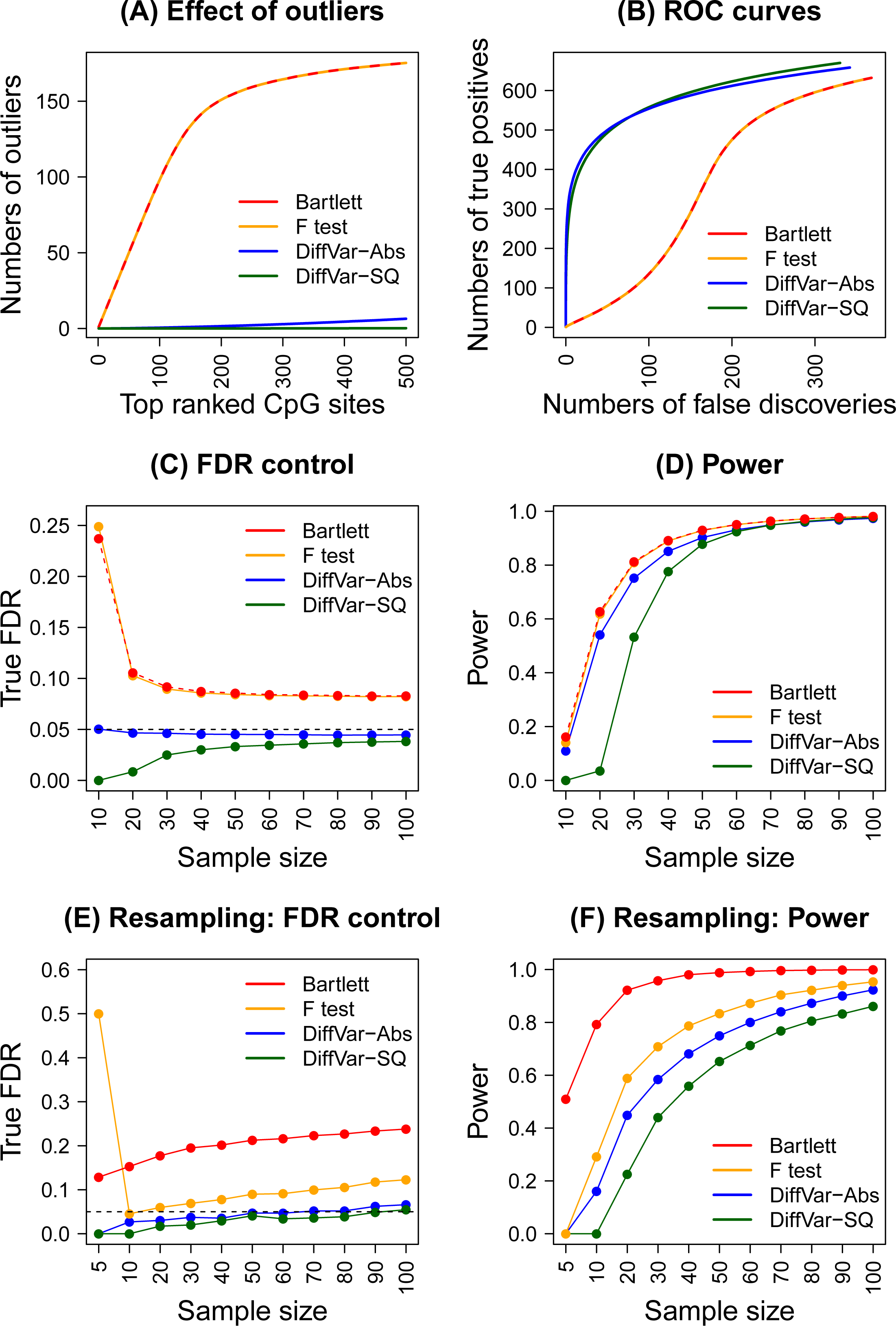
Performance of the four different methods using simulations. Results shown in plots (A-D) are averaged over 1000 simulated datasets and results shown in plots (E-F) are averaged over 100 resampled datasets at each sample size. Plot (A) shows the cumulative numbers of differentially variable CpG probes containing outlier observations when ranking by the four different methods. There is no differentially variable CpGs simulated but 200 outliers are included in the data. Plot (B) shows ROC curves when 200 outliers and 1000 differentially variable CpGs are present in the simulated data. Plot (C) shows the control of the false discovery rate (FDR) of the four methods at a 5% nominal FDR cut-off (horizontal dashed black line) over ten different sample sizes. This simulation contains 1000 CpGs which are roughly five times more variable in Group 2 compared with Group 1. Plot (D) shows the power to detect differentially variable features at ten different sample sizes when Group 2 is roughly five times more variable than Group 1. Plot (E) shows the control of the FDR using resampled kidney cancer datasets at 11 different sample sizes. Plot (F) shows power to detect differentially variable features using resampled kidney cancer datasets at 11 different sample sizes.

**Table 2:** **Comparing the type I error rate for the four different methods in the presence of 200 outliers.** Median type I error rates are reported for 1000 simulations with no differentially variable or differentially expressed features, but with 200 outliers incorporated in the data. The standard deviation with which the error rate is estimated ranges from approximately 0.0003 for rates near 0.001 to 0.003 for rates near 0.1 with the Bartlett and F test more variable than DiffVar at nominal p-values of 0.001 (sd approximately 0.002 compared to 0.0003) and 0.01 (sd approximately 0.0018 compared to 0.0010).

|  | **Nominal P-value** |  |  |  |
| --- | --- | --- | --- | --- |
| **Test method** | **0.001** | **0.01** | **0.05** | **0.1** |
| Bartlett | 0.01379 | 0.02601 | 0.06781 | 0.1179 |
| F test | 0.01467 | 0.02663 | 0.06767 | 0.1172 |
| DiffVar (abs) | 0.0009701 | 0.01006 | 0.05085 | 0.1019 |
| DiffVar (sq) | 0.0005373 | 0.008198 | 0.04880 | 0.1011 |

#### DiffVar has fewer false positives in the presence of both outliers and truly differentially variable features

To add complexity to the simulations we incorporated 1000 differentially variable features in the simulated data in addition to the 200 outliers. The variability of the 1000 CpGs in the second group was roughly twice that of the variability in the first group (see ‘Materials and Methods’). Figure 2B shows Receiver Operating Characteristic (ROC) curves for the four methods. It is clear that the outliers severely affect the F and Bartlett tests’ ability to detect the true differentially variable features. The DiffVar tests always detect more true positives and fewer false positives than the F and Bartlett tests, with very little difference between absolute or squared deviations. The dip in the curves corresponding to the F and Bartlett tests show that the outliers are ranked above truly differentially variable CpGs. DiffVar correctly ranks truly differentially variable features at the top of the list and is not affected by outliers.

#### Sample size considerations for testing differential variability

In order to accurately estimate the group variances, larger samples sizes are needed than for accurately estimating group means. We investigated a range of sample sizes that would enable reliable detection of differentially variable features. Detecting differential variability depends on sample size as well as how large the variability in one group is compared to the other. We assessed the effect of sample size for DiffVar in two ways. Firstly, through the use of simulations, and secondly, by subsetting the kidney cancer dataset which contains 283 cancer tumours and 160 normal samples.

Initially, we simulated 50,000 CpGs with 200 outliers and 5000 CpGs sites differentially variable between two groups with sample sizes of 10, 20, 30, 40, 50, 60, 70, 80, 90 and 100. We simulated three scenarios with the 5000 differentially variable CpGs having twice, five times and ten times the variability in group two compared to group one, and generated 1000 such datasets for each scenario. Total numbers of differentially variable features were assessed at 5% false discovery rate cut-off for the four different methods, with the numbers of false discoveries and true discoveries recorded for each dataset.

In addition, we used real data from the kidney cancer versus normal tissue dataset to investigate the effect of sample size. We took a conservative approach to determining the “true” differentially variable CpGs by taking the intersection of the significant CpGs at 1% FDR using DiffVar (abs), the F test and the Bartlett test when analysing the full dataset. We randomly sampled 5, 10, 20, 30, 40, 50, 60, 70, 80, 90 and 100 samples for each group and re-analysed the subsetted data in order to see how many “true” differentially variable features were recovered at 5% FDR. To estimate the false discovery rate, we classed any significant differentially variable CpG not in the list of “true” differentially variable CpGs as a false discovery. We repeated the sampling procedure 100 times for each distinct sample size.

The results for the simulated and real data are displayed in Figure 2C, D, E, F and Supplementary Figure 1. Across all simulation scenarios a similar pattern emerges. While the F and Bartlett test tend to have greater power to detect true differentially variable CpGs, this comes at a price of more false discoveries, particularly at small sample sizes (Figure 2C, D, E, F and Supplementary Figure 1). DiffVar (abs) and DiffVar (sq) control the false discovery rate correctly at all samples sizes, however DiffVar (sq) is conservative for sample sizes below 40 (Figure 2C, E and Supplementary Figure 1A, C). For data where the variability in group two is roughly twice that of group one, larger sample sizes are needed to detect differentially variable CpGs (Supplementary Figure 1B). For data where the variability in group two is roughly ten times that of group one, sample sizes as low as n=20 show more than 80% power to detect differential variability (Supplementary Figure 1D).

A similar story emerges when looking at the kidney cancer data. When n=5, the Bartlett test is the only test to recover significant CpGs in the resampled data (Figure 2F), however it shows inflated false discovery rates across all sample sizes (Figure 2E). DiffVar (abs) has almost perfect FDR control across all sample sizes, however it lacks power when n=5. A minimum sample size of 10 appears necessary to recover differentially variable CpGs (Figure 2F). For both the simulated and real datasets we find a trade-off between control of the false discovery rate and power but DiffVar (abs) gives the best compromise with a lot fewer false positives.

### Application to TCGA methylation datasets

#### Top differentially methylated and differentially variable CpG sites have little overlap

We analysed the kidney, lung adenocarcinoma and prostate cancer versus normal tissue methylation data from TCGA (see ‘Materials and Methods’ for sample details). All samples were hybridised to Illumina’s Infinium HumanMethylation450 arrays. The data was read into R using the *minfi* Bioconductor package [18], the raw intensities normalised using SWAN [39] and M values and *β* values calculated. Differential methylation was assessed by performing moderated t-statistics on the M values using the *limma* Bioconductor package. Differential variability was determined using DiffVar with absolute deviations. Significant differentially methylated CpGs were identified as those with FDR < 5% and difference in mean *β* at least 0.1. Significant differentially variable CpGs were identified as those having FDR < 5%, as well as having a variability ratio of at least five. In other words, the magnitude of the variance in one group had to be at least five times the magnitude of the variance in the second group.

Under these criteria, there were 59,271 significant differentially methylated CpGs in the kidney cancer dataset, 77,588 in the lung cancer dataset and 71,361 in the prostate cancer dataset. The proportions of these that were hyper-methylated in cancer were 47% in the kidney cancer dataset, 47% in the lung cancer dataset and 70% in the prostate cancer dataset (Figure 3A). DiffVar detected 109,529 differentially variable CpGs in the kidney cancer dataset, 146,453 in the lung cancer dataset and 22,001 in the prostate cancer dataset (Figure 3B), with the vast majority of these more variable in cancers compared with normal samples (Kidney: 99.9%, Lung: 99.8%, Prostate: 95%).

**Figure 3.**
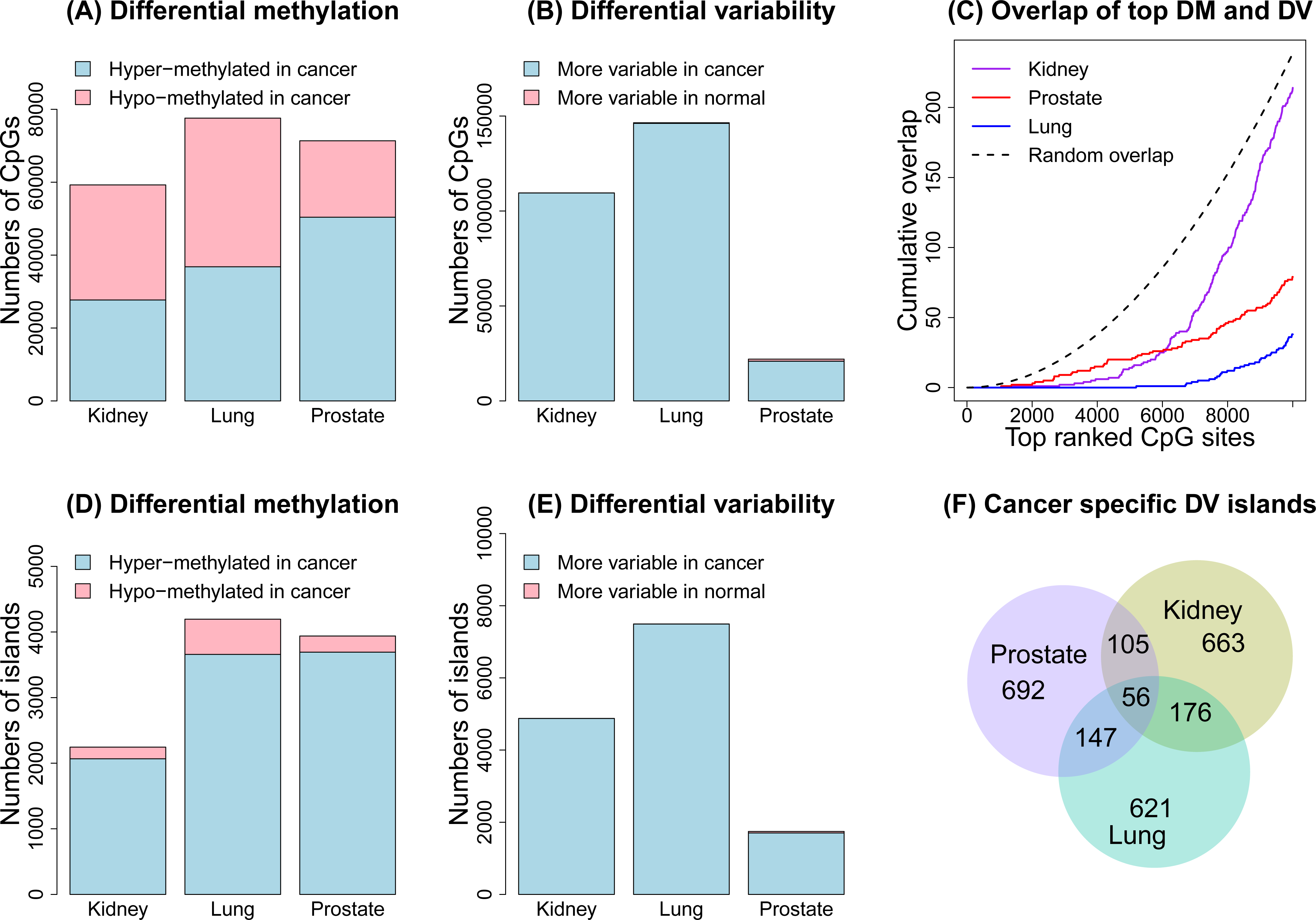
Analysis of the three TCGA datasets. The top panel (A-C) shows results of CpG site-level analysis and the bottom panel (D-F) shows results of the CpG island-level analysis. Plot (A) shows the numbers of significantly differentially methylated CpG sites for the three cancer datasets; (B) shows the numbers of significantly differentially variable CpG sites for the three cancer datasets; (C) shows the overlap of the top differentially methylated and top differentially variable CpG sites for each cancer dataset separately. The dotted line shows the median overlap profile of two sets of 10,000 randomly selected CpGs. The random sampling was repeated 1000 times. Plot (D) shows the numbers of significant differentially methylated CpG islands; (E) the numbers of significant differentially variable CpG islands; and (F) shows a proportional venn diagram of the overlap of the top differentially variable CpG islands between the three different cancers.

Restricting to only the most highly ranked significant differentially methylated and differentially variable CpGs, we found very little overlap between the top ranked 10,000 CpG sites (Figure 3C). This phenomenon was observed across all cancer datasets, with the degree of overlap between differentially methylated and differentially variable CpGs lower than expected by chance (Figure 3C). Kidney cancer had the most overlap (2%) and lung cancer had the least overlap (0.5%). This implies that the most significant differentially methylated and differentially variable CpG sites are different in the three cancer datasets analysed.

#### CpG islands are highly variable in cancer samples compared to normal tissue samples

We next investigated the genomic context of differentially variable probes. We found that the top differentially variable CpG sites were mostly in CpG islands. Furthermore there was a greater proportion of differentially variable sites in CpG islands compared to the differentially methylated CpG sites. This was consistent across the three cancer datasets (Figure 4A, Supplementary Figures 2A, 3A). An interesting observation in the kidney cancer dataset was that the CpG islands were over-represented among differentially variable CpG sites but under-represented among differentially methylated CpG sites. The lung and prostate cancer dataset had more CpG islands represented among the top ranked differentially methylated CpG sites, however there was always a greater proportion of CpG islands represented in the top differentially variable CpG sites. Hansen et al [9] found increased variability of almost all CpG sites (islands, shores and distal to islands region) in their custom made CHARM array. However, when analysing a publicly available colorectal cancer versus normal mucosa data from the Illumina Human Methylation 27K BeadChip array they found that differentially variable CpGs were over-represented in sites far from islands and shores, and under-represented in islands. This contrasts our findings, and may be due to differences in the genomic region composition of the different arrays, differences in sample size (22 matched colorectal cancer versus normal mucosa), differences in analysis strategy and potentially differences in tissue type.

**Figure 4.**
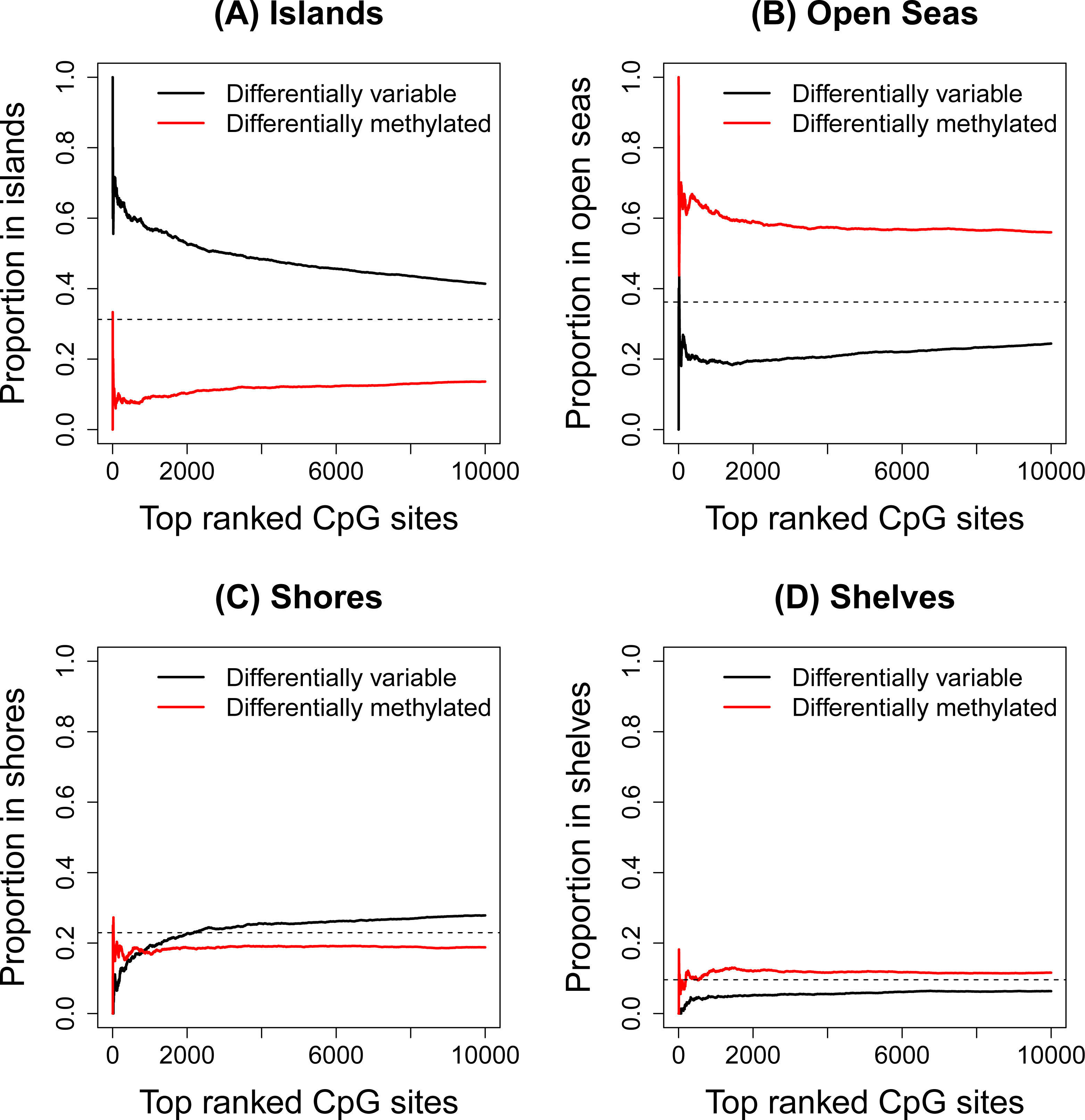
Breakdown of significant DM and DV CpG sites by genomic region in kidney dataset. Plot (A) shows the proportion of the top 10,000 ranked CpGs in islands, (B) open seas, (C) shores and (D) shelves in the kidney cancer dataset, based on UCSC annotation. The horizontal dashed line represents the overall proportion of probes for the genomic region on the array.

This led us to examine the proportion of CpG sites in shores, shelves and open seas, based on UCSC annotation (Figure 4B, C, D, Supplementary Figures 2B, C, D, 3B, C, D). A CpG that is not in an island, shore or shelf was classed as being in an open sea. In the kidney cancer dataset there is a striking difference in the proportion of differentially methylated and differentially variable CpG sites in open seas (Figure 4B), with a large number of differentially methylated CpG sites appearing in these regions. There are fewer CpG sites in shelves and shores represented on the array, hence interpretation of these regions is more difficult. In the lung and prostate cancer datasets the differences between top differentially methylated and top differentially variable CpG sites in shores, shelves and open seas is not as striking.

To gain additional insight in to the differentially variable and differentially methylated CpG sites that are in open seas, we tested whether ENCODE defined regulatory regions [40] were over-or under-represented (see ‘Materials and Methods’ for further details). Of the nine human cell types that have these regulatory regions characterised, the normal lung fibroblasts annotation is the only tissue relevant to our datasets. For the significant differentially variable CpGs in open seas in the lung cancer dataset we found that strong enhancers, weak enhancers, active promoters and weakly transcribed regions were over-represented; and heterochromatin/low signal regions were under-represented. For the significant differentially methylated CpGs in open seas, strong enhancers and active promoters were over-represented and heterochromatin/low signal regions were under-represented. The results are displayed in Supplementary Table 1.

### Top differentially variable CpG islands tend to differ between cancer types

Due to the interesting finding that the differentially variable CpG sites are over-represented in CpG islands, we proceeded with a CpG island-level analysis by averaging the intensities of the CpG sites across each CpG island. Just over 30% of the probes on the array interrogate CpG islands, with 25,744 unique islands represented. Even though between individuals the CpG sites are highly variable, within an individual, the M values of the CpG sites in a CpG island are highly correlated (Supplementary Figure 4). There were 2245 significant differentially methylated CpG islands in the kidney dataset, 4195 in the lung dataset and 3939 in the prostate dataset. The majority of the significant CpG islands were hyper-methylated in cancer compared to normal (Kidney: 92%, Lung: 87%, Prostate: 94%, Figure 3D). There were 4877 significant differentially variable CpG islands in the kidney dataset, 7495 in the lung dataset and 1745 in the prostate dataset. We found an even greater proportion of CpG islands were more variable in the cancers compared to the normal tissue (Kidney: 99.9%, Lung: 99.9%, Prostate: 98%, Figure 3E). The top 1000 differentially variable CpG islands showed a trend of being unmethylated in the normal samples and becoming methylated in the cancer samples in the Kidney dataset (Supplementary Figure 5). These CpG islands tended to differ between cancer types (Figure 3F). This agrees with the findings in Fernandez et al. [10], who show that the DNA methylation profile is tumour-type specific over 1054 cancer samples, with the tumour profiles characterised by a progressive gain of methylation in CpG island associated promoters. In our datasets, kidney and lung cancer had the most differentially variable CpG islands in common (20%), but across all three cancer datasets the overlap was minimal (approximately 5%).

### Infiltrating cells as a cause of CpG variability between cancer samples

One explanation of the increased variability seen in cancer samples is that cancer cells are frequently infiltrated with normal cells. To address this as the underlying cause of differential variability we performed a thorough analysis relating the methylation levels and tumour purity of the cancer samples in three different cancer datasets (Kidney, Lung and Uterine). We included an extra dataset from TCGA (Uterine corpus endometrial carcinoma) to get a better understanding of differentially variable CpGs that have tumour methylation signal explained by tumour purity in low (lung), medium (kidney) and high (uterine) tumour purity contexts (Figure 5A). A differential variability analysis of the Uterine dataset had 101,262 CpGs significantly more variable in the cancer samples compared to normal samples, and 773 CpGs significantly more variable in the normal samples compared to the cancer samples. The proportions of the top 10,000 differentially methylated and differentially variable CpGs in islands, open seas, shores and shelves was similar to the Kidney, Lung and Prostate cancer datasets, with a larger proportion of differentially variable CpGs in CpG islands (Supplementary Figure 6).

**Figure 5.**
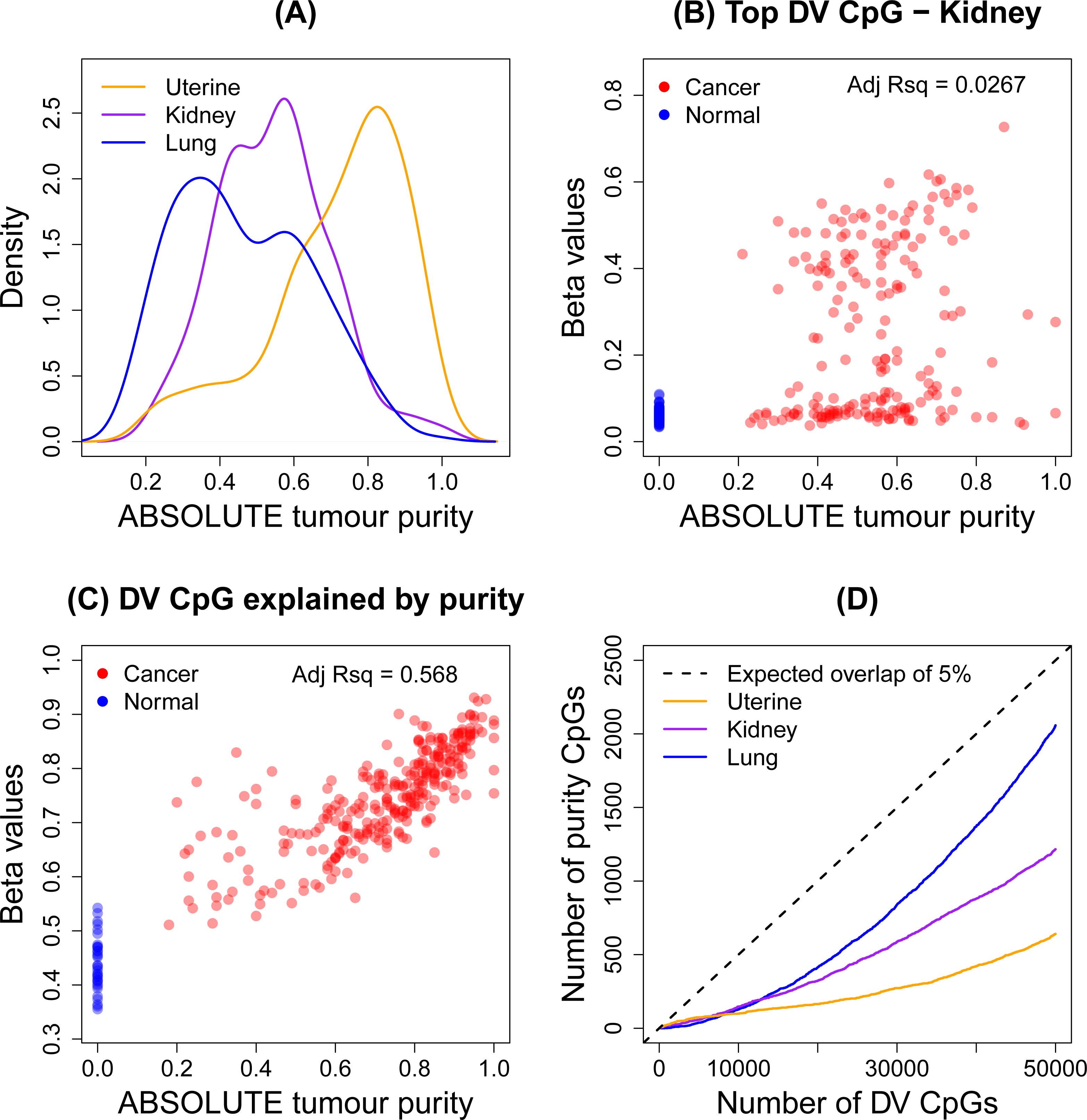
Effect of tumour purity on methylation signal in three TCGA cancer datasets. Plot (A) shows the distribution of the ABSOLUTE tumour purity estimates for Uterine, Kidney and Lung cancer datasets. Plot (B) shows the correlation of ABSOLUTE tumour purity estimates with the methylation signal for the top ranked differentially variable CpG site in the Kidney dataset. Plot(C) shows the correlation of ABSOLUTE tumour purity estimates with the methylation signal for a CpG site that is classified as differentially variable, differentially methylated and has a large R-squared value in the Uterine cancer dataset. Plot (D) shows the number of CpGs that have R-squared values of at least 10% in the top 50,000 differentially variable CpGs. The black dashed line clearly indicates that less than 5% of the top 50,000 differentially variable CpG sites are explained by tumour purity.

Carter et al. [41] have developed and applied an algorithm (ABSOLUTE) to estimate tumour purity for a number of publicly available datasets, including 11 TCGA cancer datasets, although purity estimates for the prostate cancer samples are not available. There are ABSOLUTE estimates available for 196 samples in the Kidney cancer dataset, 236 in the Lung dataset and 297 samples in the Uterine cancer dataset. Figure 5A shows a density plot of the ABSOLUTE purity estimates for three cancer datasets: Lung (median purity = 0.44), Kidney (median purity = 0.56) and Uterine (median purity = 0.76).

In order to assess the effect of tumour purity on methylation levels we fitted a linear model with methylation level as the response variable and tumour purity as the explanatory variable for the cancer samples only. The adjusted R-squared values, which can be interpreted as the proportion of the variation in the methylation signal that is explained by tumour purity, were obtained for each CpG in each cancer dataset. We classed CpGs that had an adjusted R-squared value of greater than 10% to be the CpGs that had tumour methylation signal best explained by the purity of each cancer sample. For example, for the top differentially variable CpG in the Kidney cancer dataset, Figure 5B shows that there is no strong evidence of a relationship between methylation signal and tumour purity (adjusted R-squared = 0.0267). By contrast, Figure 5C shows the relationship for a CpG (cg08395122) that is identified as differentially variable (FDR = 0.0000294, Variability Ratio = 6.14), differentially methylated (FDR = 2e-41, Δ*β* = 0.3), and has a high adjusted R-squared value (0.568) in the Uterine cancer dataset. In the case of the CpG in Figure 5C, it is clear that the variability between cancer samples is mostly explained by the tumour purity.

The numbers of CpGs on the array that had more that 10% of variability explained by purity were 29,424 in Kidney cancer, 47,353 in Lung cancer and 25,832 in Uterine cancer. However most of these were not detected as differentially variable between cancer and normal. Of the significant differentially variable CpGs, 3635 CpGs (3.3%) had at least 10% of the variation explained by tumour purity in Kidney, 14986 (10.2%) in Lung and 2745 (2.7%) in Uterine cancer. Figure 5D shows the cumulative number of CpGs with at least 10% of the variation explained by tumour purity in the top 50,000 most significant differentially variable CpGs. The Lung cancer samples, which have the lowest median tumour purity, have the largest overlap between top differentially variable CpGs and CpGs with at least 10% of the variation explained by tumour purity; while the Uterine cancer samples, which have the highest median tumour purity, show the least overlap.

### Application to aging methylation dataset

In order to study differential variability in aging we used data from Heyn et al. [2] to compare methylation in 19 centenarians to 19 newborns. We followed the same analysis strategies used for the cancer datasets to determine significant differentially methylated and differentially variable CpG sites between centenarian and newborn samples (see ‘Materials and Methods’). Of the 31,805 significant differentially methylated CpGs, 9130 (29%) were hyper-methylated in centenarians and 22,675 (71%) were hypo-methylated (Figure 6A). We also detected 34,680 significant differentially variable CpGs. Intriguingly, although perhaps unsurprisingly, 97% of these were more variable in the centenarians than in the newborns (Figure 6A).

**Figure 6.**
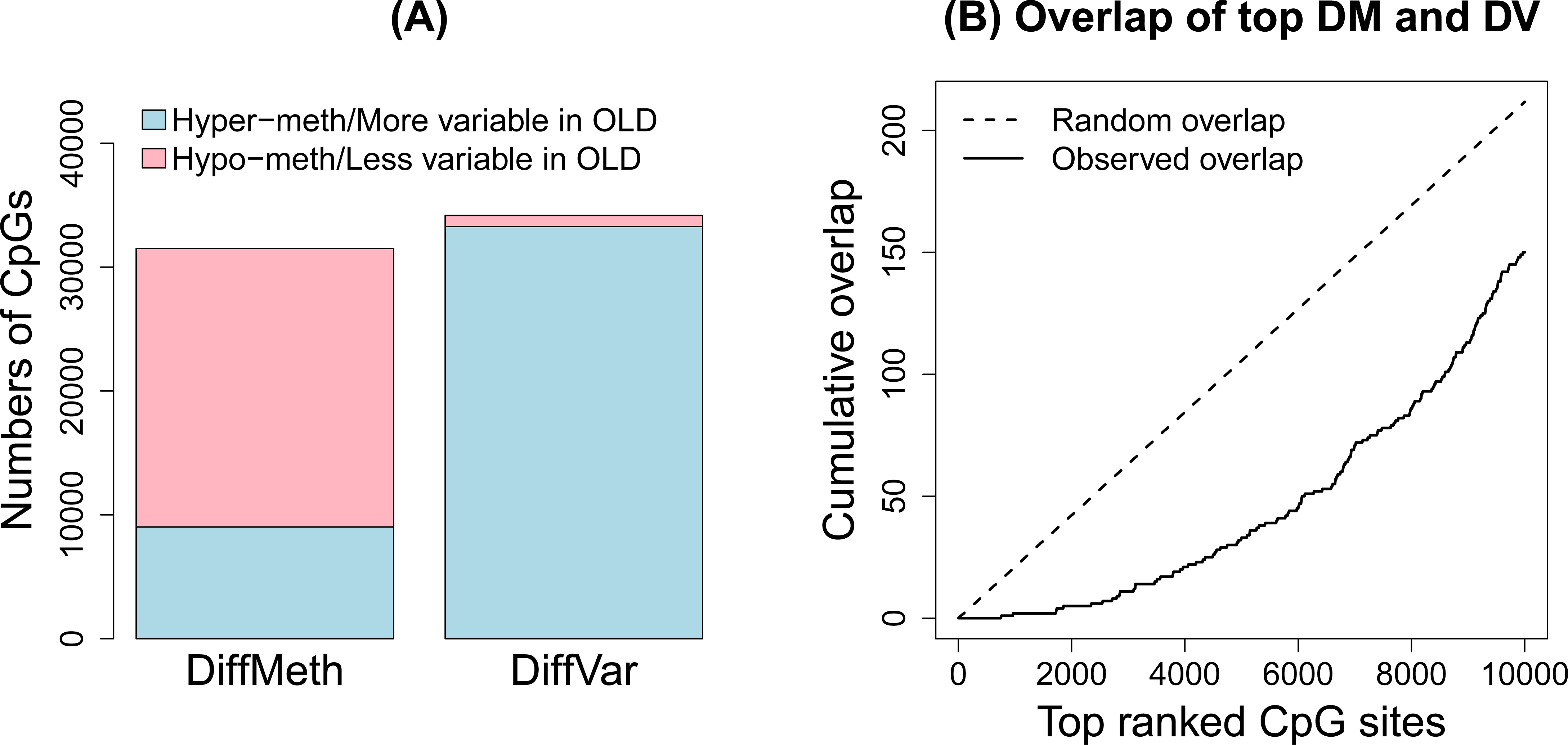
Analysis of aging dataset. Plot (A) shows the numbers of significant differentially methylated and differentially variable CpG sites and (B) shows the overlap between the top 10,000 differentially methylated and differentially variable CpG sites.

Comparing the 10,000 top ranked significant differentially methylated and top ranked 10,000 differentially variable CpGs, we observed that there was less overlap than expected by chance (Figure 6B). Of the 31,805 total significant differentially methylated CpGs and 34,680 total significant differentially variable CpGs, there were 5978 CpGs in common. Of the common CpGs, 5842 (98%) were more variable in centenarians than newborns, with 47% hypo-methylated and 53% hyper-methylated).

A closer look at the genomic composition of the significant differentially methylated and differentially variable CpGs revealed that there were more islands and shores represented in the top ranked 10,000 differentially variable CpGs compared to differentially methylated CpGs (Supplementary Figure 7A, C). There were more CpGs in open seas and shelves in the top differentially methylated CpGs than in the top differentially variable CpGs (Supplementary Figure 7B, D).

The CpGs that were significantly more variable in the centenarians compared to newborns corresponded to 7068 genes and the CpG sites that were more variable in newborns compared to centenarians corresponded to 484 genes (UCSC annotation). We performed a GOstats analysis [42] testing for over-representation of GO terms amongst the genes associated with differentially variable CpGs. For the CpGs that are more variable in centenarians compared to newborns, many GO categories relating to development were significant (Supplementary Table 2, p-value cut-off = 0.0001). Far fewer GO terms were significant when testing the genes associated with CpGs that were more variable in newborns compared to centenarians; however several GO terms related to MHC protein complex were significant, as well as a leukocyte mediated immunity GO term (Supplementary Table 2).

## DISCUSSION

In this paper we present DiffVar, a new method to detect differential variability between groups of samples. DiffVar is freely available as an R function in the *missMethyl* package (https://sites.google.com/site/oshlacklab/software/humanmethylation450). Our test is based on Levene’s z-test for variances and incorporates an empirical Bayes modelling framework to appropriately deal with high dimensional data issues. We find the test holds its size, is robust to outliers and outperforms the F and Bartlett test in terms of controlling the false discovery rate. For smaller sample sizes, the F and Bartlett test are more powerful; however this comes at a price of higher false discovery rates. DiffVar with absolute deviations represents the best compromise between controlling the FDR and power to detect differential variability.

In order to reliably estimate the variance of a group, more observations are needed than for estimating a mean. We found that for group sizes as low as 10 in the kidney cancer dataset there was adequate power to detect differential variability, with the power increasing dramatically up to group sizes of 50 arrays. However, detection of differential variability is also dependent on how large the variability in one group is relative to the other. In addition to a false discovery rate cut-off, one can specify a threshold on the variability ratio, which is the ratio of the variance in one group compared to the variance in the second group. For example, when analysing the cancer and aging datasets, in addition to the FDR cut-off of 5%, we required a variability ratio of at least five between the two groups for a CpG to be called significantly differentially variable.

We applied our differential variability testing method to several cancer data sets from TCGA as well as a publicly available aging dataset. For the cancer datasets, we found that a large proportion of sites showed differential variability and, as expected, were more variable in cancer compared with normal tissue. For the aging dataset, we found that a large proportion of CpG sites were more variable in centenarians than newborns. Interestingly, for the top ranked CpGs in both the cancer and aging datasets, the overlap between differentially variable and differentially methylated CpGs was less than expected by chance.

In all cancer datasets we showed that CpG islands were enriched for differential variability. This is important because methylation of CpG islands in cancer cells is known to contribute to gene silencing [43]. CpG sites that are differentially variable may reflect a loss of epigenetic control that could be contributing to tumourigenesis. Another hypothesis is that differentially variable CpG sites capture the heterogeneity between patients whose tumours arose due to the disruption or activation of different biological pathways. For some cancer patients, the methylation status of a differentially variable CpG site is similar to the methylation status in a normal cohort, while for other patients the same CpG site shows aberrant methylation. This type of analysis has the potential to be used as a starting point to identify clusters of patients whose tumours arose via similar molecular mechanisms. For the cancer datasets analysed in this paper, we found that only a small proportion of differentially variable CpGs could be explained by tumour purity.

In this paper we have focused on DNA methylation data that has been generated using Illumina’s Infinium HumanMethylation450 BeadChip. DiffVar can also be applied to DNA methylation sequencing data or any set of *β* or M values from CpG sites or regions. The DiffVar function will transform a matrix of *β* values into M values by applying a logit transformation. In general, the DiffVar method for testing differential variability can be applied to any ‘omics data which uses the *limma* pipeline for analysis, such as microarray expression data or RNA-seq data. For RNA-Seq data, DiffVar will perform a *voom* transformation [44] before testing for differential variability. The DiffVar modelling framework has the added benefit of giving the user access to other tools in the *limma* package, for example, testing relative to a threshold (TREAT [45]), as well as gene set testing functions ROAST [46] and CAMERA [47].

## MATERIALS AND METHODS

### Illumina Infinium HumanMethylation450 beadchip

The Illumina Infinium HumanMethylation450 beadchip allows for the simultaneous measurement of 482,421 CpG sites. There are two types of probes: Infinium I, which were previously used on the Infinium HumanMethylation27 array and constitute 135,501 probes; and Infinium II, which make up the rest of the probes. The Infinium I design has two probes which measure the methylated and unmethylated state respectively, whereas the Infinium II design has a single probe which can detect whether a CpG site is methylated or unmethylated. A thorough description of the array design is available in Maksimovic et al. [39]. The resulting data is similar to two-colour microarray gene expression data; there is an intensity measurement for both the methylated and unmethylated channel for each CpG site. The main difference is that the dynamic range for expression data is different to the dynamic range for methylation data. SWAN normalisation can be performed on the Infinium I and Infinium II probes within each array to reduce the technical biases inherent in the probe design before statistical analysis [39]. Once normalisation has been performed, *β* values for each CpG site can be computed as the ratio of the methylated intensity versus the combined methylated plus unmethylated intensity. Du et al. [34] recommend the use of M values for statistical analysis, which are calculated as the log_2_-ratio of methylated versus unmethylated intensities. A small offset can be added to the numerator and denominator to stabilise the M values. M values with an offset of 100 to both the numerator and denominator are used in all statistical analyses reported in this paper.

### Datasets

Throughout the paper we demonstrate our method with publicly available datasets from The Cancer Genome Atlas (TCGA) as well as an aging dataset, all of which have methylation profiled using the Illumina HumanMethylation450 arrays. In all TCGA datasets used, the cancer samples are from solid tumours and the normal samples are from solid normal tissue. The raw data in the form of idat files (called “Level 1” data on the TCGA website) for the “Methylation450” platform were downloaded from the TCGA data portal and sample annotation was downloaded from the Biospecimen Metadata Browser by specifying “Analyte” as the element and “DNA” as the analyte. The clear cell kidney carcinoma versus normal tissue dataset has 160 normal samples and 283 cancer samples. The lung adenocarcinoma dataset has 427 cancer samples and 31 normal samples. The prostate adenocarcinoma has 194 cancer samples and 49 normal samples. The uterine corpus endometrial carcinoma dataset had 423 cancer samples and 34 normal samples. The specific sample names and other sample information can be found in Supplementary Tables 3, 4, 5 and 6. For the aging dataset, methylation profiles were generated from 19 healthy male centenarian peripheral blood samples and 19 male newborn umbilical cord blood samples. The data is available for download from the Gene Expression Omnibus (series accession number GSE30870). The data was read into R from raw idat files using the *minfi* package. For the cancer datasets, CpG sites on the X and Y chromosomes were filtered out. An additional filter was applied to all datasets where CpG sites that had a detection p-value of greater than 0.01 in one or more samples were excluded from further analysis. This resulted in 445,378 CpG sites for the kidney cancer analysis, 419,031 CpG sites for the lung cancer analysis, 448,145 CpG sites for the prostate cancer analysis, 421,795 CpG sites for the uterine cancer analysis and 483,615 CpG sites for the aging analysis. The raw intensities were SWAN normalised, and finally *β* and M values extracted. Statistical analysis was performed on the M values. Differentially methylated CpG sites were determined using moderated t statistics from the *limma* package. Significant differentially methylated CpGs were defined as having false discovery rate (FDR) of less than 5% and a difference in mean beta level of at least 0.1 between the two groups. Differential variability was assessed using DiffVar with absolute deviations. Significant differentially variable CpGs were defined as having FDR less than 5% and a ratio of group variances of at least 5. Testing for enrichment of GO categories in the aging dataset was performed using the R *GOstats* package [42]. The R code for the analysis is available in Additional file 1. The tables of significant differentially methylated and differentially variable CpGs are available from the authors on request.

### Simulations

To check the performance of our method we generated datasets where the truth is known in order to gain insight into Type I error rate and false discovery rate control. Data was generated under a hierarchical model whereby the variance for each CpG was first sampled from an inverse chi-square distribution and the M value for each CpG was sampled from a normal distribution with variance equal to the simulated variance. A two group problem with 50 arrays in each group and ten thousand CpGs were generated in this manner. One thousand datasets with no differential variability were simulated by first sampling variances for each CpG site from a scaled inverse chi-square distribution with scaling factor *d_0_ s_0_^2^* and degrees of freedom *d_0_* such that the two groups have equal variance. We chose *d_0_* = 20, which represents the scenario where the observed variances are shrunk 20/(50+20) = 0.29 towards the prior variance. In other words, more weight is placed on the observed variance during the empirical Bayes shrinkage step. We chose *s_0_^2^* = 0.64, which is slightly more variable than the hyper-parameter estimates obtained when analysing the larger TCGA cancer datasets (Kidney: 0.26, Lung: 0.11, Prostate: 0.16). The M values were generated to mimic methylation data by randomly sampling half the CpGs from a normal distribution with mean equal to −2 and variance equal to the sampled variance for the unmethylated CpGs; and the remainder representing methylated CpGs had M-values sampled from a normal distribution with mean equal to 2 and variance equal to the sampled variance. The range of M values this simulation scenario produced is shown in Supplementary Figure 8A.

Type I error rates were also assessed using the M values of the non-diseased samples from the kidney cancer dataset. One thousand datasets were generated by first randomly sampling ten thousand of the 445,378 CpGs, followed by randomly selecting 100 normal samples, of which 50 were allocated to the “normal” group and 50 were allocated to the “cancer” group. Theoretically, for each CpG the two groups should have equal variance, although additional factors such inter-individual heterogeneity will make the data more variable. The sampling strategy should preserve any correlation structure between the CpG sites. The range of M values this produced for one such dataset is shown in Supplementary Figure 8B.

To assess the impact of outliers we modified the simulations to include outliers for 200 CpG sites. For each of these 200 CpGs the M-value for one simulated sample was replaced with the maximum simulated M value over all ten thousand CpGs. We introduced differential variability for another 1000 CpGs by simulating larger variances for one of the groups by sampling from a scaled inverse chi-square distribution with scaling factor *d_0_ s_0_^2^* and degrees of freedom *d_0_*, where *d_0_ =* 20 and *s_0_^2^* = 1.5. On average, the variability of the 1000 CpGs in the second group is 1.5/0.64 = 2.34 times larger than the variability in the first group.

Differential variability was assessed using our method DiffVar, the F test for equality of variances and the Bartlett test. All three tests had two-sided p-values computed. Type I error rates and false discovery rates were calculated and averaged over the 1000 simulations in each simulation scenario.

The simulations to assess the effect of sample size were modified slightly by increasing the numbers of simulated CpGs to 50,000 and allowing 5000 CpGs to be differentially variable. Three different levels of variability were considered: group two roughly twice, five times and ten times more variable than group one. A two group problem with varying sample size (n = 10, 20, 30, 40, 50, 60, 70, 80, 90, 100) was simulated with 1000 datasets generated for each distinct sample size. The numbers of truly differentially variable CpGs recovered at a 5% FDR and the numbers of false discoveries was recorded for each sample size. Additional file 2 shows the R code for the simulations. For the sub-setting and resampling of the kidney cancer dataset we defined a set of true positives and true negatives in the following way. CpG sites that were differentially variable at 1% FDR using DiffVar (abs), the Bartlett and F test when analysing the full dataset were classed as true positives. All other CpG sites were classed as true negatives. The false discovery rate for each sub-setted dataset was estimated by counting the number of true negatives that were significantly differentially variable and dividing by the total number of significant differentially variable CpG sites at a 5% FDR cut-off. Power for each sub-setted dataset was calculated by counting the number of true positives that were significantly differentially variable at 5% FDR and dividing by the total number of true positives.

### Overlap analysis of ENCODE regulatory regions

Ernst et al. [40] used a Hidden Markov Model to segment the genome and computationally predict functional elements for nine human cell types, using ChIP-Seq data for nine factors plus input. The chromatin state segmentation annotation is available for download at the ENCODE website (http://genome.ucsc.edu/ENCODE/). The only relevant tissue where annotation is available is for normal lung fibroblasts. We determined whether CpG sites that were significantly differentially variable and in open seas (as opposed to islands, shores and shelves) lay in regions that have predicted functional elements using an intersectBed analysis in the Lung cancer dataset. We calculated the total number of each functional element represented on the array by taking all CpGs in open seas and performing an intersectBed analysis. We then calculated the probability of each functional element being over-or under-presented in the list of significant differentially variable and differentially methylated CpGs separately using a hypergeometric test. The p-values were adjusted for multiple testing using Holm’s method [48].

### Infiltrating cells analysis

ABSOLUTE tumour purity estimates for the Kidney, Lung and Uterine cancer datasets were downloaded from the Synapse website (www.synapse.org). Adjusted R-squared values were used to determine the proportion of variation in the tumour methylation signal explained by the tumour sample purity. Adjusted R-squared values were extracted from a linear model fit, regressing the methylation signal on the ABSOLUTE tumour sample purity estimates of the cancer samples for all CpG sites. This was performed in R using the *lm* function. CpG sites that had adjusted R-squared values of more than 10% were classed as CpGs that had methylation signal explained by tumour purity.

## ABBREVIATIONS

TCGA: The Cancer Genome Atlas; FDR: false discovery rate; ROC: receiver operating characteristic; DM: differential methylation; DV: differential variability.

## ACKNOWLEDGEMENTS

We thank Jovana Maksimovic who downloaded and provided R code for pre-processing the kidney cancer and aging dataset, assisted in downloading the prostate and lung cancer datasets, provided valuable advice on the analysis of methylation data, as well as providing feedback on the paper. We thank Gordon Smyth for his insightful discussion on Levene’s test. We thank Patrick Molloy for performing initial analysis related to this work. We thank Ian Majewski for providing helpful insights on the manuscript. We thank Oliver Sieber and Chris Love for helpful discussion surrounding cancer epigenetics and the biological relevance of differential variability in methylation data. The research was done with the support of NHMRC grant 1051402. AO is a career development fellow of the NHMRC 1051481.

## AUTHORS’ CONTRIBUTIONS

AO conceived the idea. BP performed the analysis and wrote the code. AO and BP wrote the paper. All authors read, edited and approved the final manuscript.

## ADDITIONAL DATA FILES

The following additional data are available with the online version of this paper. Additional file 1 (AdditionalFile1.txt) contains the R code for analysing the cancer and aging datasets. Additional file 2 (AdditionalFile2.txt) contains the R code for the simulations. Supplementary table 1 (SupplementaryTable1.xlsx) shows the results for testing over-and under-representation of ENCODE defined regulatory regions in significant DM and DV CpGs in open seas in the lung cancer dataset. Supplementary table 2 (SupplementaryTable2.xlsx) shows the GOstats analysis performed on the aging data. Supplementary table 3 (SupplementaryTable3.xlsx) shows the sample information for the kidney cancer data. Supplementary table 4 (SupplementaryTable4.xlsx) shows the sample information for the lung adenocarcinoma cancer data. Supplementary table 5 (SupplementaryTable5.xlsx) shows the sample information for the prostate cancer data. Supplementary table 6 (SupplementaryTable6.xlsx) shows the sample information for the uterine cancer data. The supplementary figures file (SupplementaryFigures.pdf) shows the additional figures referenced in the manuscript. The supplementary text (SupplementaryText.pdf) contains the statistical model details of DiffVar.

